# Bayesian non-parametric clustering of single-cell mutation profiles

**DOI:** 10.1101/2020.01.15.907345

**Authors:** Nico Borgsmüller, Jose Bonet, Francesco Marass, Abel Gonzalez-Perez, Nuria Lopez-Bigas, Niko Beerenwinkel

**Author notes:** These authors contributed equally to this work (Nico Borgsmüller, Jose Bonet).

## Abstract

The high resolution of single-cell DNA sequencing (scDNA-seq) offers great potential to resolve intra-tumor heterogeneity by distinguishing clonal populations based on their mutation profiles. However, the increasing size of scDNA-seq data sets and technical limitations, such as high error rates and a large proportion of missing values, complicate this task and limit the applicability of existing methods. Here we introduce BnpC, a novel non-parametric method to cluster individual cells into clones and infer their genotypes based on their noisy mutation profiles. BnpC employs a Dirichlet process mixture model coupled with a Markov chain Monte Carlo sampling scheme, including a modified split-merge move and a novel posterior estimator to predict clones and genotypes. We benchmarked our method comprehensively against state-of-the-art methods on simulated data using various data sizes, and applied it to three cancer scDNA-seq data sets. On simulated data, BnpC compared favorably against current methods in terms of accuracy, runtime, and scalability. Its inferred genotypes were the most accurate, and it was the only method able to run and produce results on data sets with 10,000 cells. On tumor scDNA-seq data, BnpC was able to identify clonal populations missed by the original cluster analysis but supported by supplementary experimental data. With ever growing scDNA-seq data sets, scalable and accurate methods such as BnpC will become increasingly relevant, not only to resolve intra-tumor heterogeneity but also as a pre-processing step to reduce data size. BnpC is freely available under MIT license at https://github.com/cbg-ethz/BnpC.

## Introduction

Cancer is an evolutionary process characterized by the accumulation of mutations that drive tumor initiation, progression, and treatment resistance [1]. The interplay between variation and selection ultimately leads to multiple coexisting cell populations (clones) that differ in their genotypes [2, 3]. This genomic heterogeneity, also known as intra-tumor heterogeneity (ITH), poses major challenges for cancer treatment as parts of the tumor may already be therapy-resistant [4, 5]. Therefore, it is beneficial to identify the clonal composition of the tumor and to adapt the treatment accordingly.

Recent advances in the field of single-cell DNA sequencing (scDNA-seq) have led to new insights into cancer evolution and ITH. Examples include the detection of rare subclones in breast cancer patients [6], the identification of novel treatment resistance clones in glioblastomas [7], and major advancements in the reconstruction of cancer evolution [8]. Compared to bulk sequencing, scDNA-seq offers the possibility to directly access clonal genotypes at the cellular level and to more easily detect branching in clonal evolution. However, scDNA-seq data tends to be very noisy. Experimental procedures, such as DNA amplification, but also analytic ones like alignment and mutation calling introduce a large fraction of errors in the data as well as missing values [9]. Errors can be either missed true mutations, namely false negatives (FN), or mutations not present in a cell but falsely reported, namely false positives (FP). Characteristic of scDNA-seq data are high FN rates, arising from the technical failure to measure both alleles at a mutated locus, and a large fraction of missing values, resulting from non-uniform coverage and drop-outs.

Generic clustering algorithms, such as partitioning or density-based methods, do not account for scDNA-seq characteristics and are therefore unsuitable for this type of data. Hence, various methods were recently introduced tailored to single-cell mutation profiles, i.e. the absence or presence of called mutations in each cell. These approaches differ in their main objective, model choice, and inference scheme. The majority of them focuses on resolving the phylogenetic relationship among cells and in doing so can also provide clusters and genotypes [10–14]. Currently, the only method focusing entirely on clustering and genotyping is SCG [15], which uses a parametric model and applies mean field variational inference (VI) to learn genotypes and the clonal composition. Alternatively, the centroid-basted clustering approach celluloid [16] adapts k-modes with a novel dissimilarity for scDNA-seq data but does not provide any genotyping. The probabilistic frameworks BitPhylogeny [17] and SiCloneFit [18], and the nested effects model OncoNEM [19] jointly cluster cells into clones and infer their phylogenetic relations. Despite these successes, the growing size of scDNA-seq data sets challenges the scalability of these methods, compromising their accuracy and efficiency. Especially the inference of phylogenetic relations is a computational expensive task that scales poorly with data size due to difficulties in the tree search.

Here, we introduce BnpC, a fully Bayesian method to analyse large-scale scDNA-seq data sets and to accurately determine the clonal composition and genotypes, handling noisy data and an unknown number of clones non-parametrically. We benchmark our approach against state-of-the-art methods on simulated data using various data sizes and demonstrate that BnpC outperforms current methods in terms of accuracy, runtime, and scalability. We also reanalyze published scDNA-seq data, and with our method not only manage to recapitulate the original results, but we also resolve populations that in the original publications were detected only with additional data or after manual pre-processing steps.

## 2 Methods

### 2.1 Model

BnpC takes as input a binary matrix with missing values ***X*** = (*x_ij_*) ∈ {0, 1, −}^*N×M*^ of *N* cells and *M* mutations, where 0 indicates the absence of a mutation, 1 its presence, and – a missing value (Fig. 1 A). We assume that the *N* cells were sampled from an unknown number *K* of clones, each with a distinct mutation profile *θ_k_* ∈ [0, 1]*^M^*, coming from a prior distribution *G*_0_. The probabilities of observing a FP or FN in the cell data are given by the parameters *α* and *β*, respectively, with prior distributions as stated in Figure 1 B. The assignment of cells to clones is represented by a vector ***c***, where *c_i_* = *k* is the assignment of cell *i* to clone *k*. To model the cell assignments ***c***, we use a Chinese Restaurant Process (CRP) [20]. The CRP is a probability distribution over partitions of the natural numbers, which in our model are cell assignments. Because each partition is a possible way of clustering cells, the CRP serves as a prior distribution for grouping cells into clones. The concentration parameter *α*_0_ of the CRP determines the probability of assigning a cell to a novel clone.

**Fig. 1:**
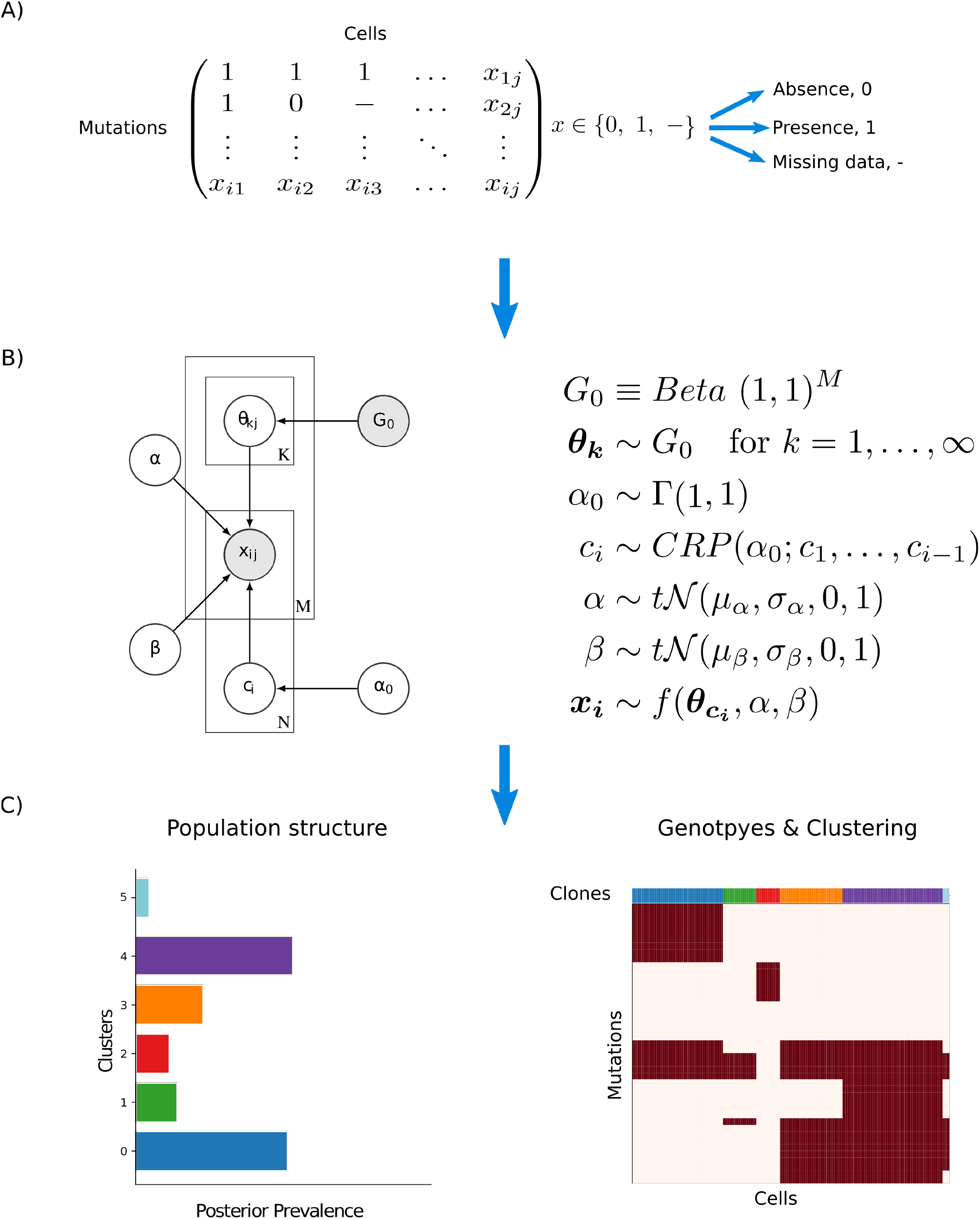
BnpC model overview. **A)** The model’s input is a binary mutation matrix, where each row represents a mutation and each column represents a single cell. Possible values are 0, indicating the absence of a mutation, 1, indicating the presence of a mutation, and missing values. **B)** BnpC’s probabilistic graphical model. The binary input data *X*, consisting of *N* cells and *M* clones, contains a fraction of FP and FN entries, indicated by *α* and *β*, respectively. *G*_0_ is a base distribution over the genotypes *θ* of an infinite number of clones. *c* is the assignment of cells to the clones, sampled from a CRP with concentration parameter *α*_0_, and *f*(·) is the model’s likelihood. Shaded nodes represent observed or fixed values, while the values of unshaded nodes are learned using MCMC. **C)** BnpC predicts clonal composition, corresponding genotypes, and the population structure.

With the parameters described above, we can formulate the likelihood of BnpC as

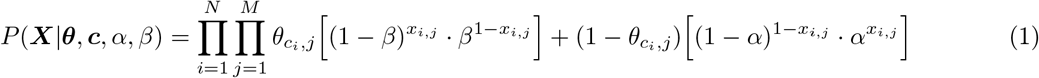

where the first term accounts for the presence of a mutation in a clone and the observation of a true positive or FN, and the second term accounts for the absence of a mutation in a clone and the observation of a true negative or a FP. Missing values are skipped.

The full posterior distribution over the latent variables factorizes as

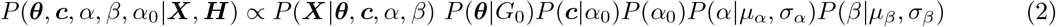

where *μ_α_, σ_α_, μ_β_*, and *σ_β_* are fixed hyperparameters.

### 2.2 Inference

As the posterior distribution (eq. 2) is not analytically tractable, we employ a Markov chain Monte Carlo (MCMC) sampling scheme, in particular a generalised Gibbs sampler, to obtain samples from the posterior distribution. Cluster parameters and error rates are updated via Metropolis-Hastings moves; the concentration parameter *α*_0_ is learned as described in [21]; cell assignments are updated with Gibbs sampling and a modified non-conjugate split-merge move [22, 23].

We modified the split-merge move introduced by Jain and Neal [24] to increase the probability of merging small clones. We first choose which move to perform. For a split move, two cells are drawn from a clone selected proportionally to its size; for a merge move, two cells are drawn from different clones, themselves selected in a manner inversely proportional to their size. This increases the probability of merging spurious clones. To account for these changes, the proposals’ ratio in Metropolis-Hastings update is modified as follows. For a split move, we introduce the ratio

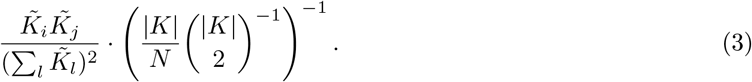

The second term describes the probability of choosing clone *K* (of size |*K*|) according to its size, and choosing two cells (*i* and *j*) from it. After the split, let *K_i_* and *K_j_* denote the two different clones to which *i* and *j* belong. Let 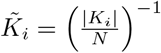 represent the inverse clone size of the clone with cell *i*. The first term in Eq. 3 denotes the probability of choosing the clone with cells *i* and *j* to reverse the split move.

Similarly, for a merge move we extend the Metropolis-Hastings ratio with the following factor:

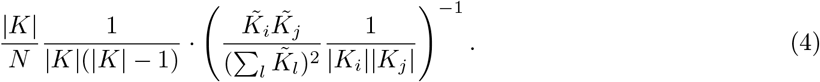

Here, the second term accounts for choosing two distinct clones in a manner inversely proportional to their size, and then two cells *i* and *j* uniformly from each clone. The first term undoes the merge move by selecting the merged clone K according to its size, and selecting cells *i* and *j* from it.

To assess convergence, we implemented an updated version of the Gelman-Rubin diagnostic [25] and compared posterior means of scalar quantities from multiple chains with random starting positions (Fig. S4). BnpC can be run for a given number of MCMC iterations, with a given time limit, or until the convergence diagnostics drop below a given threshold.

### 2.3 Estimators

Downstream analyses and interpretation generally require a single clustering and genotypes for all cells, and thus the posterior samples obtained with our model need to be summarized. To provide an estimate of the inferred clones we followed an approach inspired by MPEAR [26]. We first inferred the number of clones as the rounded posterior mean from our non-parametric model and then performed agglomerative clustering on the posterior co-occurrence matrix. The genotypes were subsequently inferred independently for each clone from a selected subset of posterior samples. For each clone, we selected posterior samples based on two criteria: *(1)* all cells assigned to the clone are clustered together; *(2)* no other cell is clustered with these cells. If no posterior sample fulfills both criteria, we selected samples fulfilling only the first criterion. The final genotype is the rounded mean over the cluster parameters from the selected posterior samples. While biased, this estimator performed well in practice. We evaluated it against MPEAR, maximum likelihood (ML), and maximum *a posteriori* (MAP). All of the mentioned estimators are implemented in BnpC and the user can choose which one to use.

## 3 Results

### 3.1 Benchmarking on simulated data

We generated 180 data sets varying the numbers of cells (1250, 2500, 5000, 10000), mutations (200, 350, 500), and clones (25, 50, 75) to assess BnpCs scalability, run time, and performance. Each combination was simulated five times with fixed FN, FP, and missing value rates at 0.3, 0.001, and 0.2, respectively. The underlying phylogeny was also fixed (minimal trunk size of 0.1 and mutation rate of 0.25). A description of the simulation process is provided in the Supplementary Material (Section Simulations). All algorithms were run four times per data set with different seeds. Clustering accuracy was evaluated using the V-Measure [27], where high values correlate with more accurate clusterings. Genotyping accuracy was measured as 1 – Hamming distance/(#cells · #mutations) between the predicted cellular genotypes and the true ones (higher values correlate with more accurate genotypes).

We benchmarked BnpC against SCG [15] with binarized output and SiCloneFit [18]. Celluloid clustering with the silhouette method for clone number determination was excluded as it does not provide genotypes and performed poorly (Fig. S8). Methods aiming to resolve phylogenetic relations were excluded as they only provide genotypes directly, while the inference of clones from phylogenetic trees is a non-trivial task. We also excluded BitPhylogeny and OncoNEM, which jointly infer clones and their phylogenetic relations, as both were previously shown to produce less accurate results than SCG and SiCloneFit [15, 18]. BnpC was run for 0.25, 0.5, 1, 2, 4, and 8 hours. SCG was run with a large number of iterations (> 10 ×10^6^) to reach convergence in every run, which SCG measures by the improvement over iterations. We were only able to run SiCloneFit for 10 steps on the data set with the smallest number of cells, where its runtime already exceeded 48 hours (Fig 2). Therefore, we excluded SiCloneFit from the benchmarks on larger data sets. Algorithms were run on a high performance computing cluster, each algorithm ran on a single core with a maximum of 128 GiB memory and 2.4 GHz CPU.

**Fig. 2:**
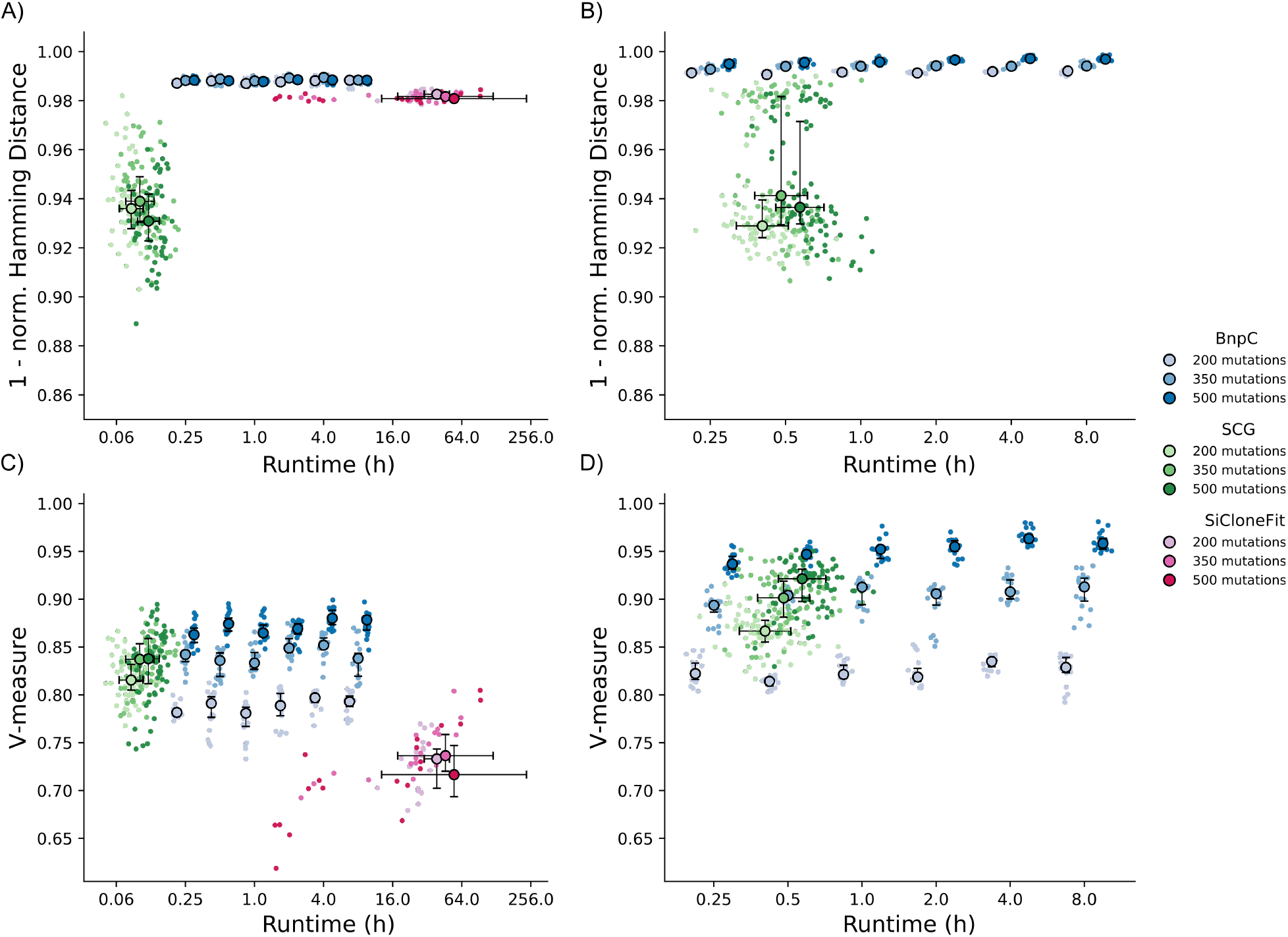
Clustering accuracy measured by the V-Measure and genotyping accuracy measured by 1 – Hamming distance/#cells · #mutations of BnpC, SCG and SiCloneFit. **A, B)** Genotyping accuracy. **C, D)** Clustering accuracy. Measured on 30 simulated data sets with six different sizes (five data sets per size): **A, C)** 1250 cells and **B, D)** 2500 cells, and 200, 350, and 500 mutations. All data sets contained 50 distinct clones, a FN rate of 30%, a FP rate of 0.1%, and a missing value fraction of 20%. Algorithms were run 4 times per data set.

On the data sets with 1250 cells and 50 clones (Fig. 2 A, C), BnpC predicted genotypes best in all cases and clones in two-third of the cases. Additionally, BnpCs predictions showed the least variance. SCG had the shortest runtime but also inferred genotypes the least accurate. On all data sets, SCGs clustering scored higher than SiCloneFits and on data sets with 200 mutations also higher than BnpCs. SiCloneFits average runtime was ≥ 48 hours and varied highly between runs. On the 2500 cell data sets (Fig. 2 B, D), the accuracy of predictions followed the same trend. BnpCs runtime to reach its highest scores was comparable to SCGs.

On the 5000 cell data sets, BnpC predicted genotypes better than SCG in all cases and achieved its maximum clustering and genotyping accuracy at shorter runtimes than SCG (Fig. S2 A, C). Moreover, we were not able to run SCG on the largest data sets of 5000 cells and 500 mutations due to insufficient memory allocation. On the 10000 cell data sets (Fig. S2 B, D), BnpC was the only method able to obtain results. Its predictions did not improve after 2 h and were ≥ 0.98 for the genotyping and ≥ 0.83 for the clustering.

Additionally, we benchmarked BnpC on smaller data sets of 50 mutations, 200 cells, and 10 clones, varying in evolutionary histories, error rates and missing value fraction (Fig. S8). In general, trends of the previous benchmarking on larger data sets recurred. BnpC predicted genotypes best except for high FN and missing value rates, where SiCloneFit performed best. On average, SCG predicted clusters best but genotypes worst, especially at a large fraction of FP. SiCloneFit predicted genotypes and clones less accurate than BnpC, except for the previously mentioned cases. We observed the same trends independent of the simulated evolutionary histories. Unsurprisingly, the simulation of different evolutionary histories showed that frequent and early branching events, resulting in clones with highly diverse mutation profiles, lead to a higher clustering accuracy of all methods than linear evolution and late branching events.

To evaluate the scalability of BnpC, we investigated the runtime per MCMC step according to the data size (Fig. S9). On the benchmarking data sets, BnpCs runtime increased linearly with data size, independently of the number of clones in the data.

We also evaluated the performance of our novel estimator on simulated data. The quality of the inferred clones varied only slightly between the estimators, not favoring any in particular (Fig. S11). To evaluate the genotype inference, we compared our estimator against ML and MAP since MPEAR only produces clusters. In all cases, our novel estimator achieved better genotyping than the point estimators, independently of the error rates and fraction of missing values (Fig. S12). All estimators showed the same trends regarding varying error rates and missing values, with higher rates resulting in less accurate inferences, and vice versa.

### 3.2 BnpC performance on tumor scDNA-seq

We analyzed the sequencing data of five patients with childhood leukemia [28], one high-grade serous ovarian cancer (HGSOC) patient [29] and two colorectal cancer (CRC) patients [30].

#### Acute Lymphoblastic Leukemia

We reanalyzed scDNA-seq data of five Acute Lymphoblastic Leukemia (ALL) patients [28]. The data contains between 16 and 105 mutations and between 96 and 143 cells per patient. Gawad *et al*. used a combination of a multivariate Bernoulli model and the Jaccard distance to predict the clonal composition and to infer genotypes. Inferred genotypes and clones by Gawad *et al*. as well as the ones inferred by BnpC are displayed in Fig. S10. Genotypes and clones predicted by BnpC are largely in accordance with those previously determined. BnpC predicted some additional clones of small size. BnpC predictions were of partly higher resolution. Specifically for patient 4, BnpC was able to detect an additional clone (orange) differing from the closest clone by five mutations (Fig. 3 A). The identification of this particular clone results in a different and more accurate evolutionary pattern, as a common ancestor for the two tumor branches is obtained (Fig. 3 B). Gawad *et al*. confirmed the existence of this additional clone in their subsequent analysis by incorporating copy number data. These findings show that our approach is sensitive to small clones and able to recover biological meaningful results.

**Fig. 3:**
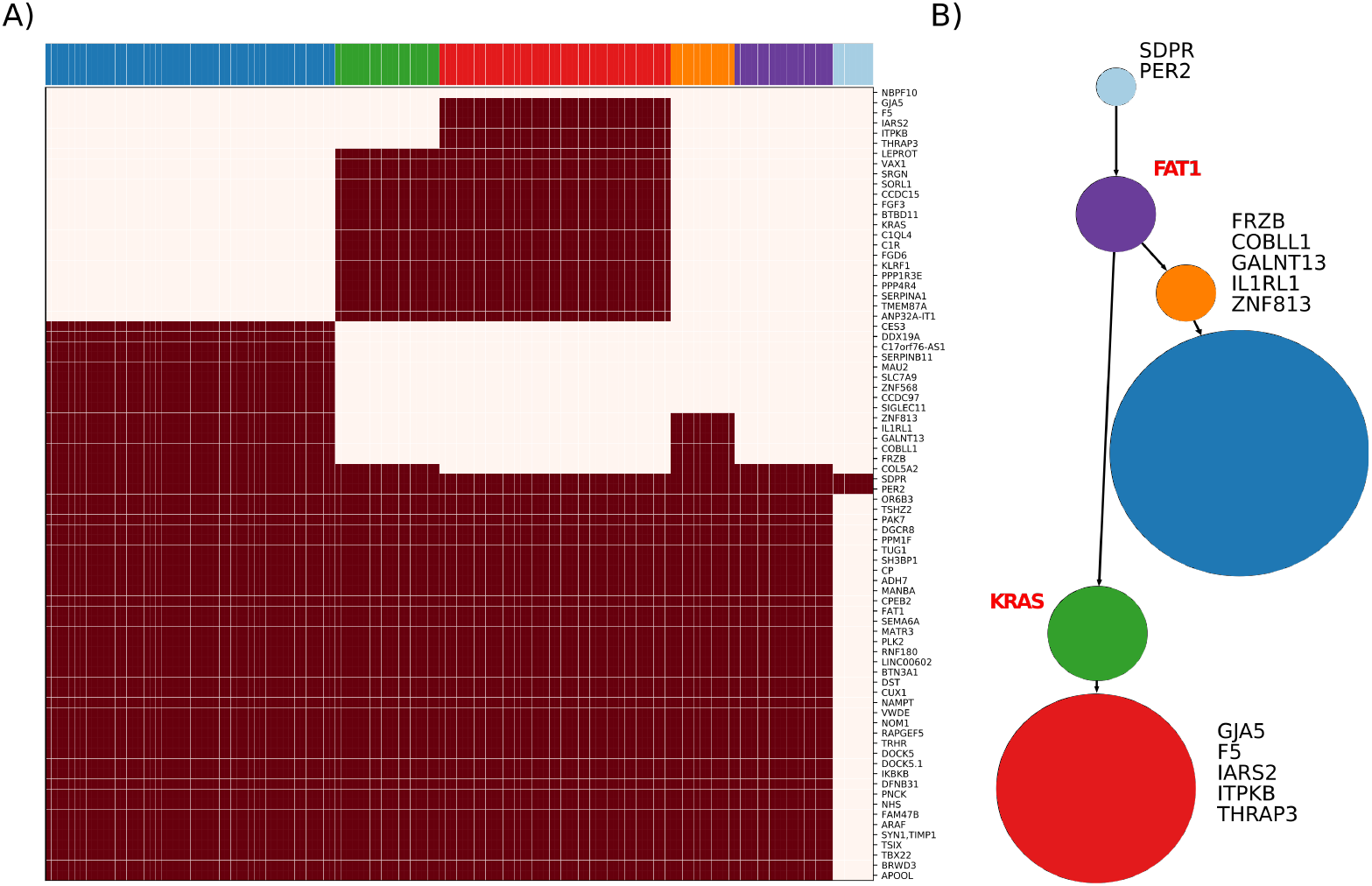
**A)** Clones and genotypes inferred by BnpC for patient 4 of the Gawad data set. Heatmap depicts absence (white) or presence (red) of mutations for every mutation (row) in every cell (column). **B)** Resulting minimum spanning tree from the clonal genotypes as obtained in Gawad et al. Gene labels in the tree determine either mutations leading to a new clone (black) or known ALL driver genes (red). Node size corresponds with the clonal size.

#### High-Grade Serous Ovarian Cancer

The HGSOC data of patient 9 from the McPherson data set [29] was obtained by whole-genome sequencing of five samples taken from three tumor sites: left ovary, right ovary, and omentum. The data consists of 420 cells, 43 SNVs, and five breakpoints. We compared our predictions to the results obtained by Roth *et al*. using SCG. Their initial clustering analysis identified a normal population and eight tumor clones, of which they filtered out three clones due to a high fraction of missing values in the corresponding cells (mean ≥ 20% SNV events missing per cell).

BnpC was able to produce the same findings as SCG [15] without applying any additional filtering step (Fig. S13). By excluding the three clusters, 28 cells which represent 7% of the patient data were discarded.

The clonal prevalence shows differences between the two samples coming from the left ovary (LOv) (Fig. S13 B). Populations within one of the two samples (LOv2) contain the amplification in ERBB2, while the other (LOv1) does not. These populations harboring the amplification correspond to clones 0 (purple) and 1 (orange). Knowing that the primary site of the tumor was in the left ovary and that all other clones carry this amplification, our findings are in accordance with Roth *et al*.

#### Colorectal Cancer

Patients CRC0827 and CRC0907 from Wu *et al*. [30] were collected by single-cell Whole Exome Sequencing on CRC tissue samples (C1 and C2) and matched normal tissue (N). Additional samples from normal polyp (NP, CRC0907) and adenomatous polyp tissue (AP, CRC0827) were sequenced for the analysis. While BnpC recapitulated the results for patient CRC0827, we identified an additional clone for patient CRC0907 (green clone in Fig. 4). This new clone suggests another step in the clonal evolution of the tumor. For patient CRC0907, Wu *et al*. identified two tumor clones harboring somatic mutations. They subsequently analyzed a subset of functionally related mutations to CRC development and separated them into unique clonal (detected by bulk sequencing) and unique subclonal (not detected). The results obtained from BnpC allows us to further classify the unique subclonal mutations into early subclonal (contained in the green clone) or late subclonal events (only present in the blue clone). Therefore, our method suggests an early mutation of LAMA4 compared to the other subclonal mutations annotated by Wu *et al*. (PDE3A, AB13BP, LHCGR, and CFHR5), which are only present in the later evolved population. Besides, we observed one of their annotated unique clonal mutations (STXBP1) to be present only in the blue clone, suggesting a later acquisition of the mutation. In summary, these results indicate that BnpC can give new insights into the evolution of the tumor and the order in which mutations are acquired by better identifying the clonal composition within tumor samples.

**Fig. 4:**
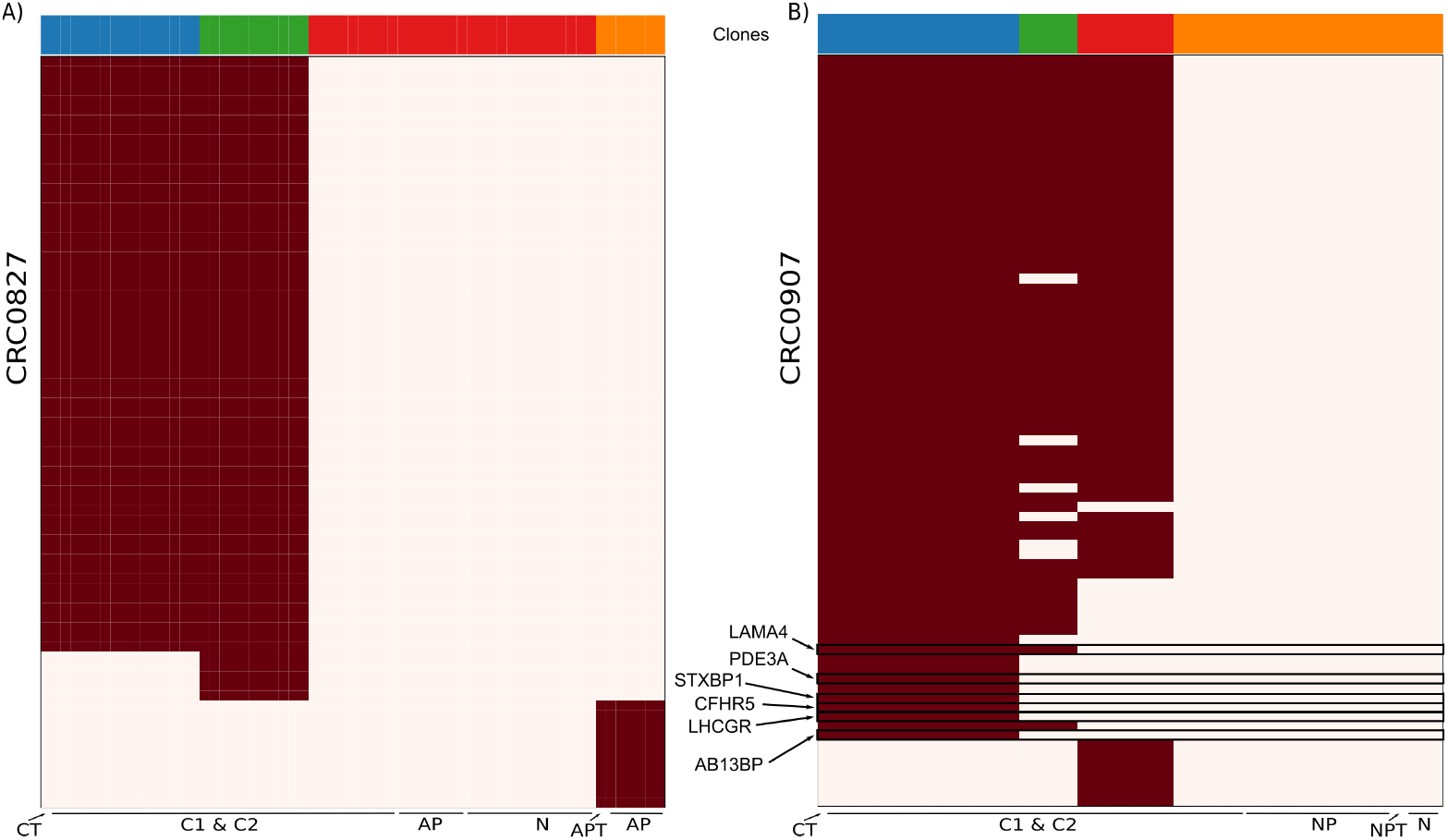
Inferred clones and genotypes by BnpC for patients CRC0827 **A)** and CRC0907 **B)** of the Wu data set. Heatmap depicts absence (white) or presence (red) of mutations for every mutation (row) in every cell (column).

## 4 Discussion

The identification of the heterogeneous tumor composition and the clonal genotypes is potentially advantageous for cancer treatment. ScDNA-seq provides the opportunity to resolve ITH in greater detail and to detect rare clones, despite experimental protocols still producing a high fraction of FN and missing events. We have introduced the novel non-parametric probabilistic method BnpC, specially designed for accurate and scalable clustering and genotyping of large-scale scDNA-seq data. The method implements a modification of a non-conjugate split-merge move and employs a novel estimator inferring from the posterior distribution for more accurate genotype predictions.

We compared our method with the state-of-the-art methods SCG and SiCloneFit on simulated and biological data. On synthetic data, BnpCs recovered genotypes with the smallest error in all cases, and inferred clusters most precisely in the majority of scenarios. In some cases, SCG’s clustering performance was better, but its genotypes were less accurate. Inferring imprecise genotypes has a larger effect on downstream analyses than suboptimal clusters, and may mislead biological interpretation.

On biological data, our method did not only reproduce previous findings for three different data sets but identified additional clones not detected in the original analysis but confirmed by additional data in Patient 4 from Gawad *et al*. These findings highlight that more accurate analytic methods can identify signal and lead to biological conclusions which can otherwise only be drawn from additional experimental data. Additionally, we demonstrated that BnpC is able to recapitulate previous results for patient 9 from McPherson *et al*. without the manual pre-processing step conducted in the original analysis. This is of special interest, for example, for an automated analysis pipeline, where one tries to minimize manual intervention without losing accuracy.

A limitation of the BnpC model is the absence of a phylogenetic structure on cells. The information given by the mutation order could be used to correct errors in the data or to infer missing values. It is possible that this is why SiCloneFit is more robust to noise in the data on small data sets. However, approaches that use a tree structure are computationally expensive and scale poorly with data size, as seen in the benchmarking. The trade-off between accuracy, runtime, and possible optimizations needs to be investigated further. A possible extension of BnpC could be the incorporation of doublets, two single cells pooled and measured together during sequencing. Currently, doublets would be reported as separate clones. Identifying and handling them explicitly as doublets could improve the clustering and genotyping, especially of the clones corresponding with the two doublet cells.

In summary, the non-parametric nature and sampling scheme of our model results in robust clonal composition and genotype predictions in reasonable computational time. Besides their relevance for personalized treatment, the inferred clusters and genotypes can be used to reduce data size significantly, thereby facilitating downstream analyses. Therefore, a potential application of BnpC on large-scale data sets, would be as a pre-processing step for the inference of phylogenetic trees or large-scale scDNA-seq data. Additionally, not assuming a tree-structure makes our method applicable to other fields. For example, our method could be used for the analysis of methylation profiles or the analysis of microbiome data, where the input matrix indicates the presence or absence of species in samples.

As scDNA-seq data size continues to grow due to biotechnological progress, scalable and accurate inference, as provided by BnpC, will be increasingly relevant. Thus far, BnpC is the only method to predict clonal genotypes accurately on large data sets within a reasonable time. It was the only method capable of running and obtaining results for the 10,000 cell data set in reasonable time and its runtime scaled linearly with data size.

## 5 Software availability

BnpC was implemented in Python 3.7 and is freely available under MIT license at: https://github.com/cbg-ethz/BnpC.

## 6 Supplementary Material

Additional figures, a description of the simulation scheme, and a comparison of the probabilistic models of used methods.

## 7 Author Contribution

N.Bo., J.B., and F.M. designed the study. N.Bo., J.B., and F.M. developed the methodology. N.Bo. and J.B. implemented the methodology. N.Bo. and J.B. performed analyses. N.Be., A.GP., and N.LB. supervised the study. All authors drafted the manuscript and approved the final version.

## 8 Acknowledgement

This work was funded by ITN-CONTRA EU grant H2020 MSCA-ITN-2017-766030. IRB Barcelona is a recipient of a Severo Ochoa Centre of Excellence Award from the Spanish Ministry of Economy and Competitiveness (MINECO; Government of Spain) and is supported by CERCA (Generalitat de Catalunya). Part of this work was supported by the European Research Council, ERC Synergy Grant 609883 (to N.B.)

