## Supplementary material for "Bayesian non-parametric clustering of single-cell mutation profiles"

### Bayesian nonparametric clustering of single-cell mutational profiles: Supplementary Material

A Preprint

[1]Department of Biosystems Science and Engineering, ETH Zürich, 4058 Basel, Switzerland
[2]SIB, Swiss Institute of Bioinformatics, 4058 Basel, Switzerland
[3]Institute for Research in Biomedicine (IRB Barcelona), The Barcelona Institute of Science and Technology, Baldiri Reixac, 10, 08028 Barcelona, Spain
[4]Research Program on Biomedical Informatics, Universitat Pompeu Fabra, Barcelona, Catalonia, Spain
[5]Institució Catalana de Recerca i Estudis Avançats (ICREA), Barcelona, Spain
[*]Both Authors contributed equally to this study

February 28, 2020

### 1 Simulations

To simulate different evolutionary patterns as reported in recent studies [1], we generated rooted mutation trees varying in their trunk size and in the number of branching events (Fig. S1). In mutation trees, nodes represent mutations and edges represent the temporal order of mutation events. To generate mutation trees, we started from a root node (representing a normal cell) and attached mutations successively to the tree's leaf nodes. To obtain different evolutionary histories, we varied two parameters: the minimal trunk size and the branching rate. The minimal trunk sizes defined the number of sequential mutations before the first branching event occurs. After the tree reached this size, mutations were attached to a uniformly sampled leaf node. The branching rate defined the probability of a branching event, i.e. the probability of the new node being attached sequentially or in parallel, starting a new branch. In the latter case, the predecessor of the leaf node became a branching point. This process introduces a bias towards many parallel branches originating from a single branching point. By dividing the branching rate of leaf nodes by the number of parallel leaf nodes, we corrected this bias. Subsequently, we attached cells to the generated mutation tree, deduced their genotypes, added FP and FN errors, and inserted missing values according to the rates of the data set.

As it is unlikely to sequence clones from throughout the evolutionary history and to better access the clustering ability, cells were only attached to an arbitrary number $n_{cl}$ of mutations, thus, forming $n_{cl}$ distinct clones. Each leaf node formed a clone, ensuring that all mutations are represented. Remaining clones were sampled proportionally to the distance from the root. Cells were then assigned uniformly to the clones, with a minimum of one cell per clone. To avoid more leaf nodes than $n_{cl}$, we set the branching rate to 0 once the number of leaf nodes equaled $n_{cl}$.

### 2 Comparison of models and inference schemes

In this section, we briefly summarize the differences in model design and applied inference between BnpC and the other two methods compared to in our benchmarking, SiCloneFit and SCG.

#### 2.1 SiClonefit

SiCloneFit jointly infers the clonal composition, genotypes, and the phylogenetic relation between clones by employing a tree-structured infinite mixture model. The model uses a tree-structured CRP as a prior for binary (phylogenetic) trees and models the evolution of clonal genotypes with a finite-site evolutionary model adopted from SiFit [2]. In contrast, BnpC only infers the clonal composition and genotypes, and employs no evolutionary model, meaning that cluster parameters are independent of each other.

SiCloFit implements an MCMC sampling procedure based on a generalized Gibbs sampler. Cell assignments are updated via partial reversible-jump and partial Metropolis-Hastings moves, the clonal phylogeny and model of evolution via Metropolis-Hastings. Errors are updated via rejection sampling. In contrast, BnpC updates assignments via a mixture of Gibbs sampling and an adjusted split-merge move. Error rates are updated via Metropolis-Hastings as well.

**Differences summary:**

- Phylogenetic modelling with binary tree-structured CRP prior
- Model of evolution for genotypes' prior

#### 2.2 SCG

SCG employs a finite mixture model to infer the clonal composition and genotypes. To deal with the unknown number of clones, the total clone number should be set by the user to a value much larger than expected. The idea is that only fewer clones would have cells assigned to them. SCG operates on ternary data, differentiating between hetero- and homozygous mutations, and models error rates with a Dirichlet prior. In contrast, BnpC employs an infinite model, i.e. it learns the unknown number of clones, operates on binary data, treating hetero- and homozygous mutations alike, and uses a truncated normal as error rate prior.

To infer the models parameters, SCG employs mean-field variational inference (VI) and utilizes the evidence lower bound to assess convergence. In contrast, BnpC employs a MCMC sampling scheme.

**Differences summary:**

- Finite mixture model
- Definition of error rates and ternary genotypes
- Mean-field variational inference

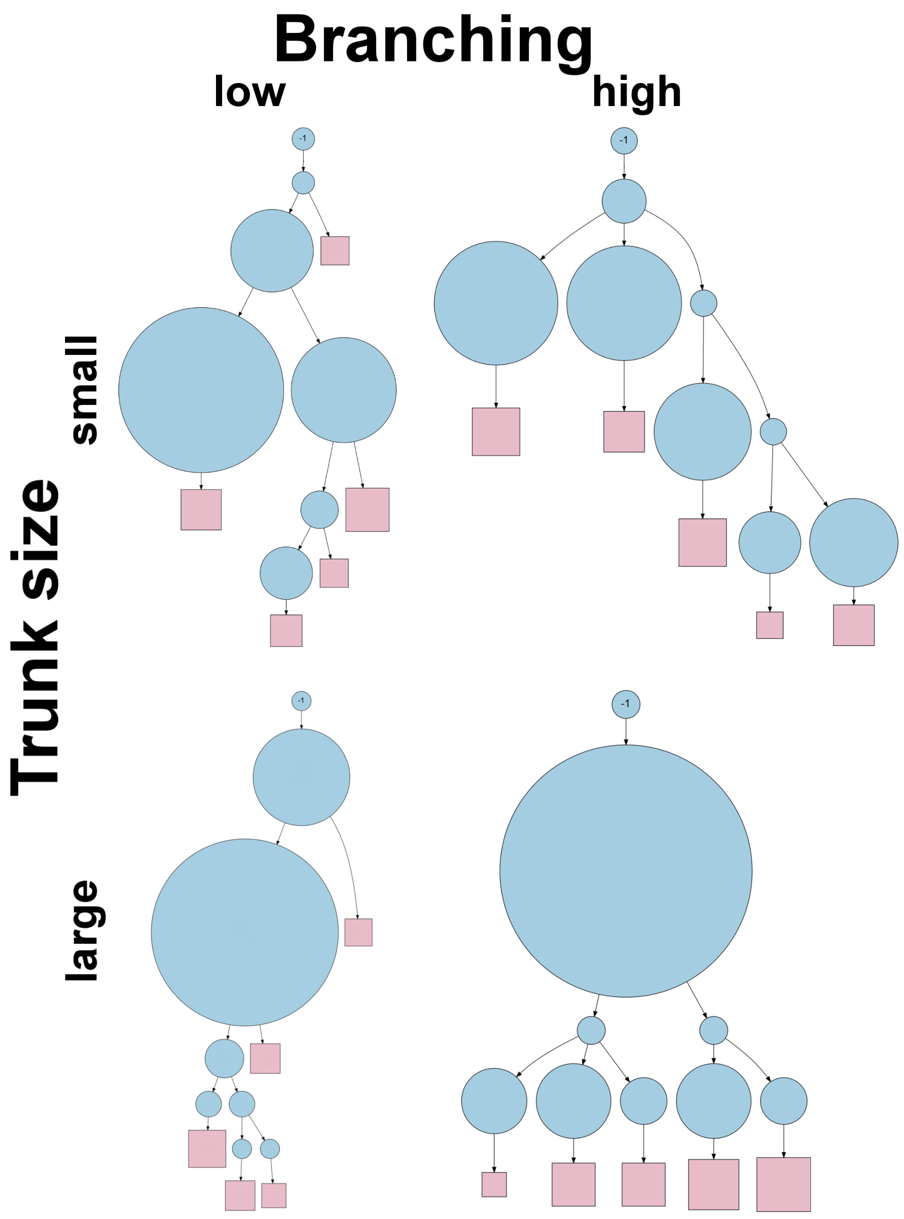


**Figure S1:** Tree topologies at different minimal trunk size and branching frequencies.

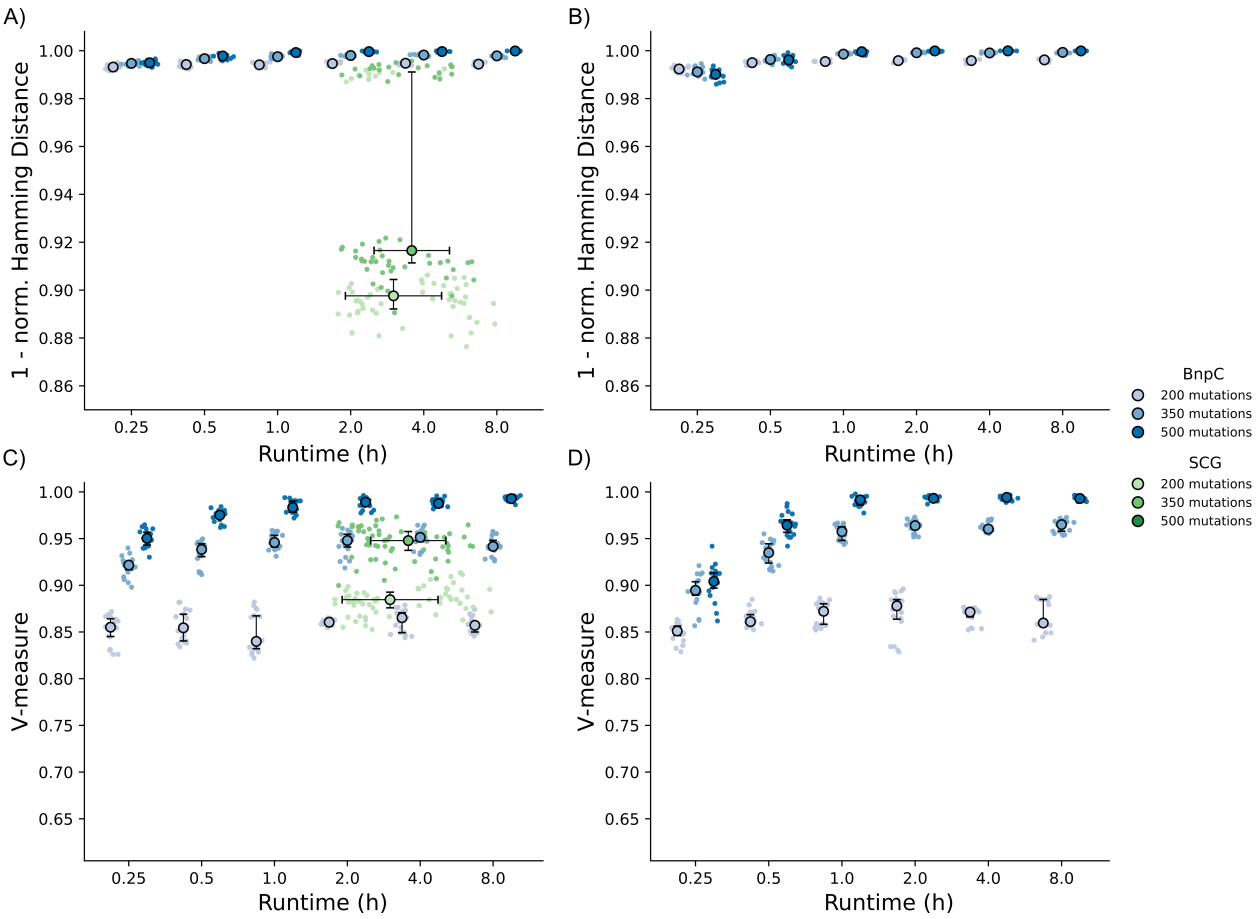


**Figure S2:** Clustering accuracy measured by the V-Measure and genotyping accuracy measured by $1 - \text{Hamming distance}/\#\text{cells} \cdot \#\text{mutations}$ of BnpC and SCG. **A, B)** Genotyping accuracy. **C, D)** Clustering accuracy. Measured on 30 simulated data sets with six different sizes (five data sets per size): **A, C)** 50,00 cells and **B, D)** 10,000 cells, and 200, 350, and 500 mutations. All data sets contained 50 distinct clones, a FN rate of 30%, a FP rate of 0.1%, and a missing value fraction of 20%. Algorithms were run 4 times per data set.

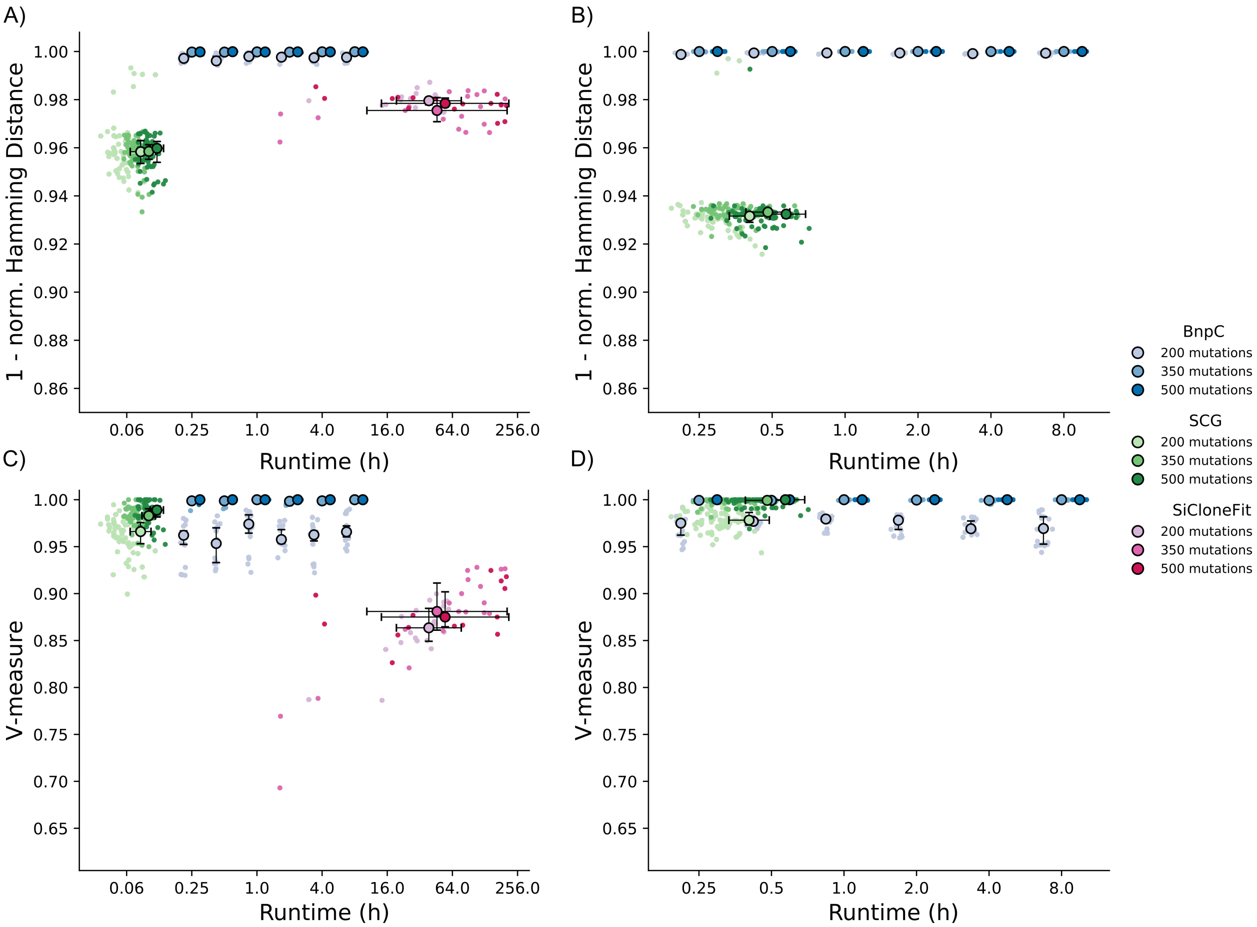


**Figure S3:** Clustering accuracy measured by the V-Measure and genotyping accuracy measured by $1 -$ Hamming distance/#cells $\cdot$ #mutations of BnpC and SCG. **A, B)** Genotyping accuracy. **C, D)** Clustering accuracy. Measured on 30 simulated data sets with six different sizes (five data sets per size): **A, C)** 1,250 cells and **B, D)** 2,500 cells, and 200, 350, and 500 mutations. All data sets contained 25 distinct clones, a FN rate of 30%, a FP rate of 0.1%, and a missing value fraction of 20%. Algorithms were run 4 times per data set.

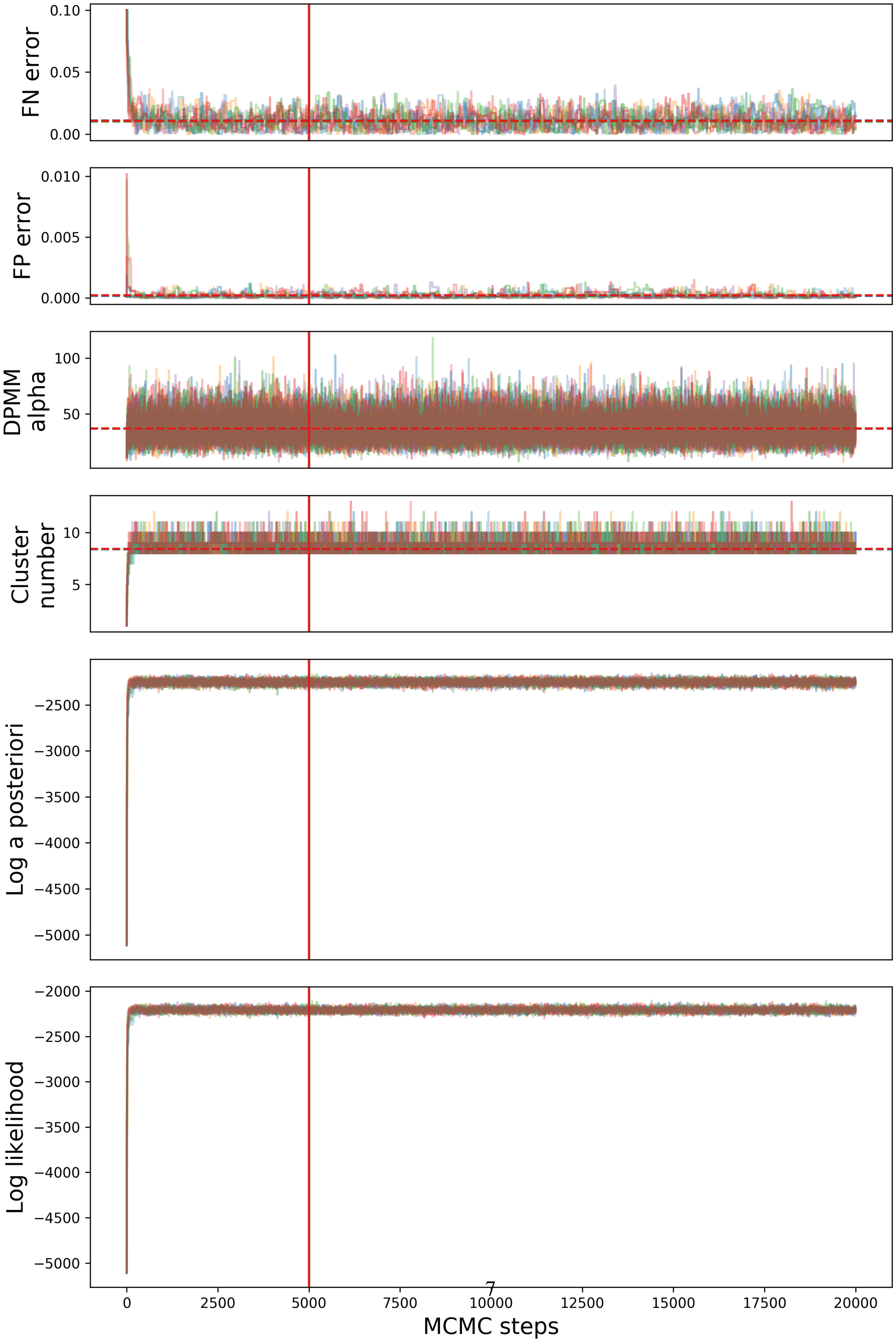

FN error
0.10
0.05
0.00
FP error
0.010
0.005
0.000
DPMM alpha
100
50
Cluster number
10
5
Log a posteriori
−2500
−3000
−3500
−4000
−4500
−5000
Log likelihood
−2000
−2500
−3000
−3500
−4000
−4500
−5000
0
2500
5000
7500
10000
12500
15000
17500
20000
MCMC steps

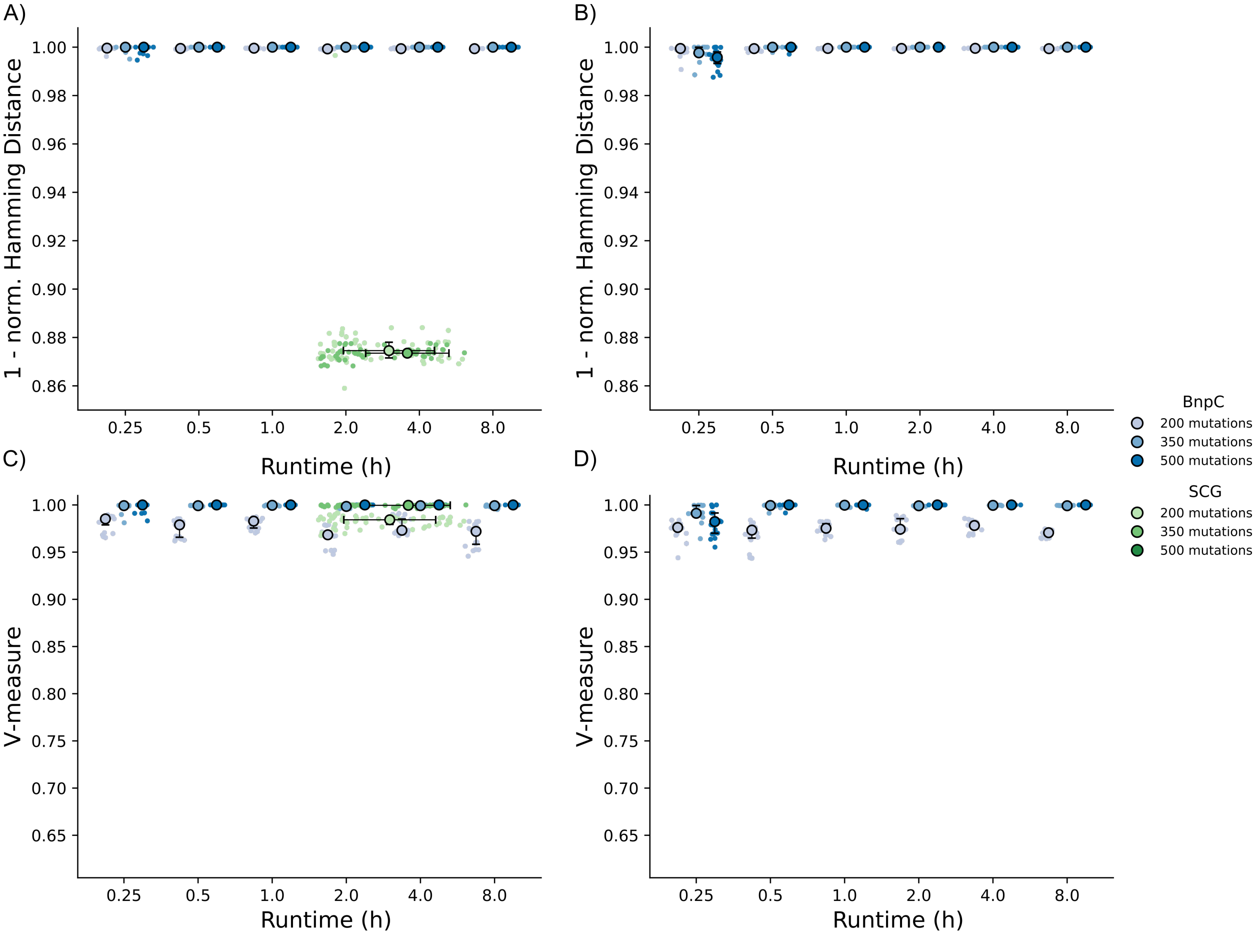


**Figure S5:** Clustering accuracy measured by the V-Measure and genotyping accuracy measured by $1 - \text{Hamming distance}/\#\text{cells} \cdot \#\text{mutations}$ of BnpC and SCG. **A, B)** Genotyping accuracy. **C, D)** Clustering accuracy. Measured on 30 simulated data sets with six different sizes (five data sets per size): **A, C)** 5,000 cells and **B, D)** 10,000 cells, and 200, 350, and 500 mutations. All data sets contained 25 distinct clones, a FN rate of 30%, a FP rate of 0.1%, and a missing value fraction of 20%. Algorithms were run 4 times per data set.

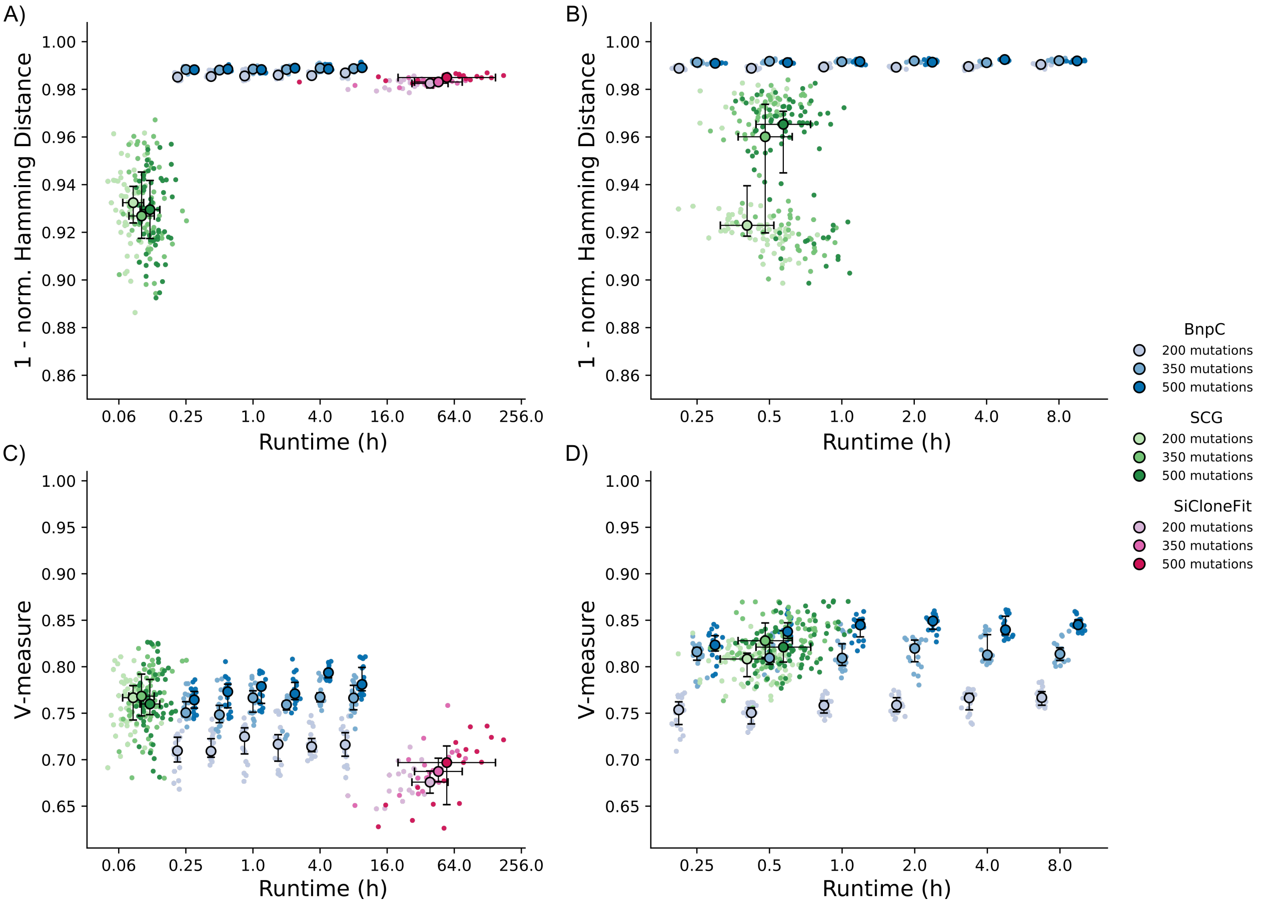


**Figure S6:** Clustering accuracy measured by the V-Measure and genotyping accuracy measured by $1 - $ Hamming distance/#cells $\cdot$ #mutations of BnpC and SCG. **A, B)** Genotyping accuracy. **C, D)** Clustering accuracy. Measured on 30 simulated data sets with six different sizes (five data sets per size): **A, C)** 1,250 cells and **B, D)** 2,500 cells, and 200, 350, and 500 mutations. All data sets contained 75 distinct clones, a FN rate of 30%, a FP rate of 0.1%, and a missing value fraction of 20%. Algorithms were run 4 times per data set.

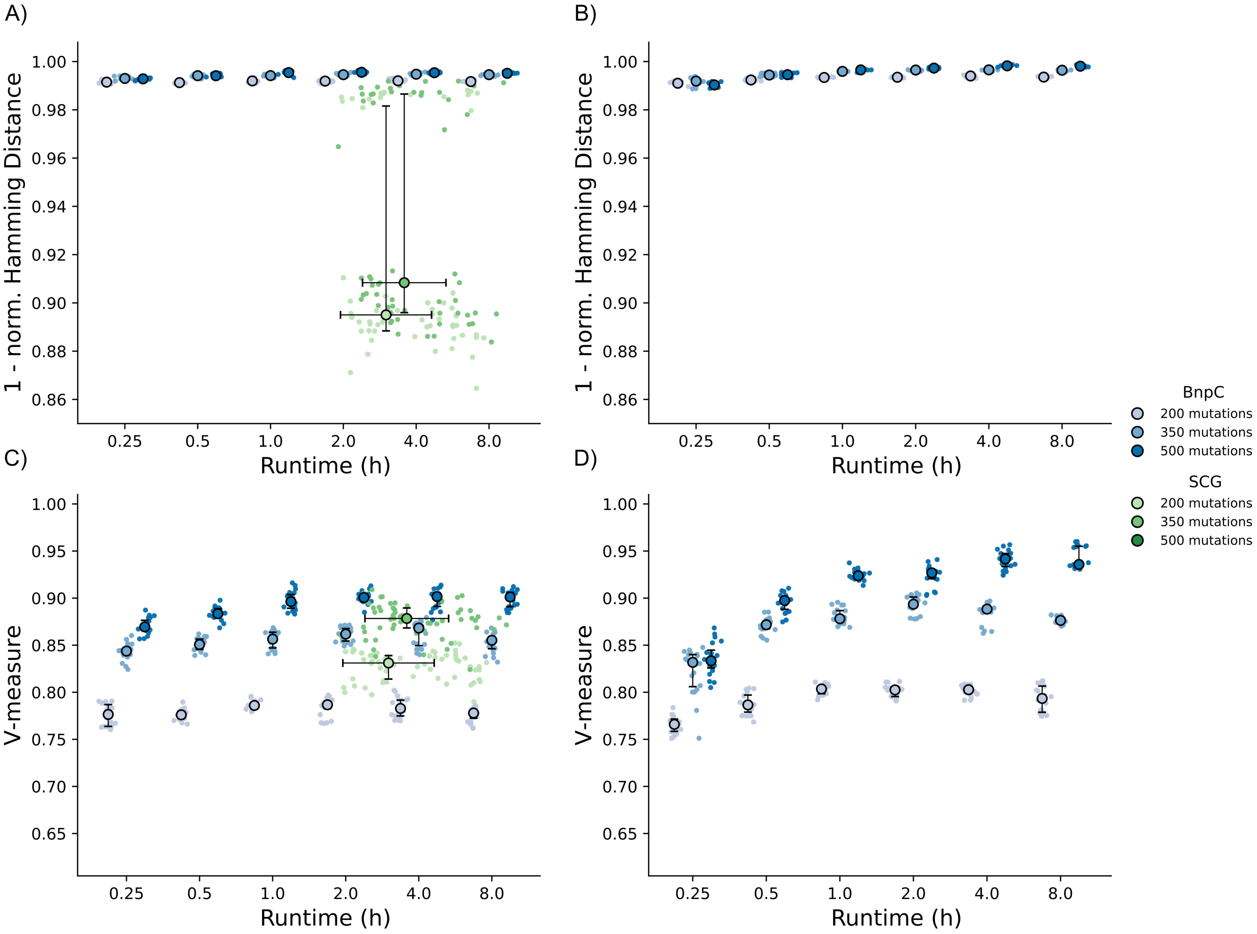


**Figure S7:** Clustering accuracy measured by the V-Measure and genotyping accuracy measured by $1 - $ Hamming distance/#cells $\cdot$ #mutations of BnpC and SCG. **A, B)** Genotyping accuracy. **C, D)** Clustering accuracy. Measured on 30 simulated data sets with six different sizes (five data sets per size): **A, C)** 5,000 cells and **B, D)** 10,000 cells, and 200, 350, and 500 mutations. All data sets contained 75 distinct clones, a FN rate of 30%, a FP rate of 0.1%, and a missing value fraction of 20%. Algorithms were run 4 times per data set.

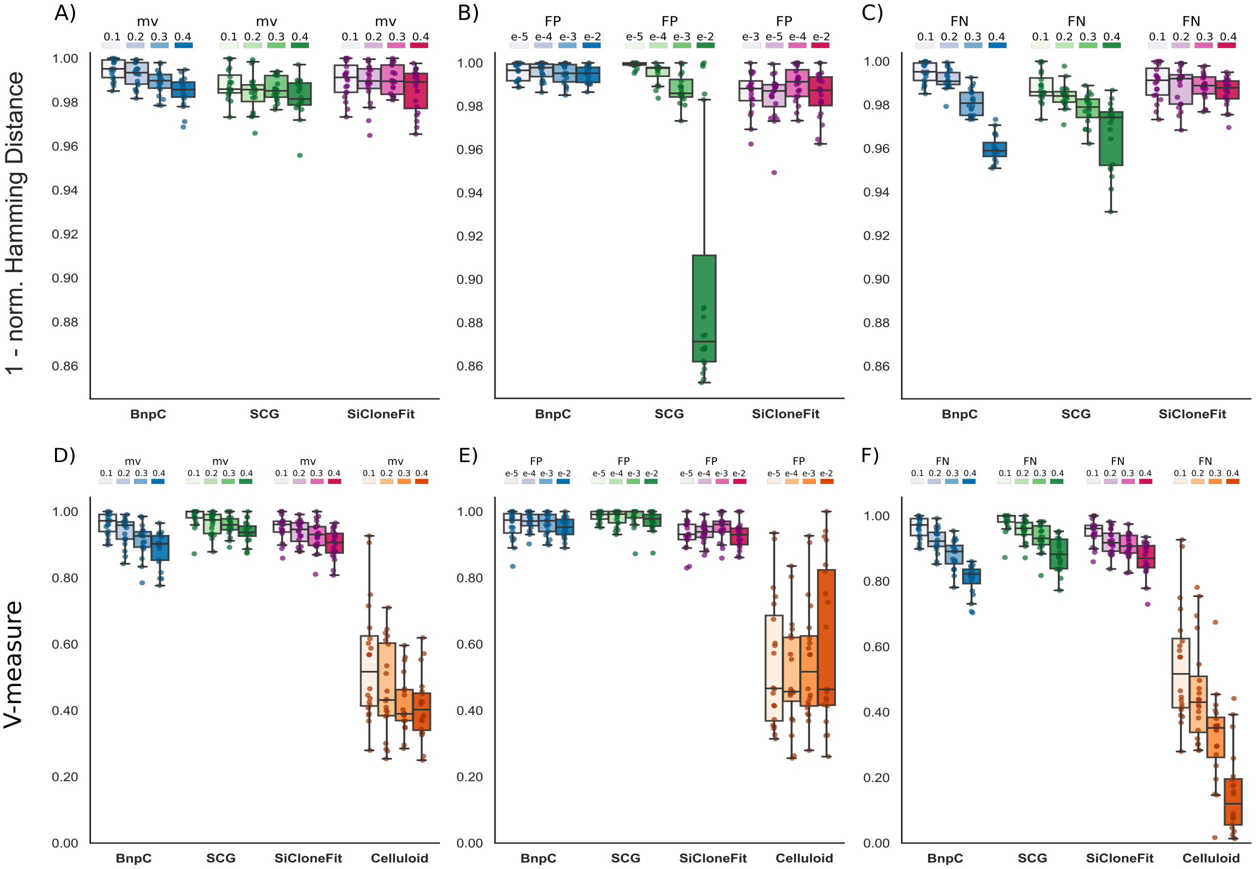


**Figure S8:** (**A, B, C**) Clustering accuracy and (**D, E, F**) genotyping error on a synthetic data set of 50 mutations, 200 cells, and 10 clusters of BnpC, SCG, SiCloneFit, and celluloid clustering. **A)** 10% FN rate, 0.1% FP rate, and a variable rate of missing values (mv). **B)** 10% missing values, 10% FN rate, and a variable FP rate. **C)** 10% missing values, 0.1% FP rate, and a variable FN rate. **D)** 10% FN rate, 0.1% FP rate, and a variable rate of missing values (mv). **E)** 10% FN rate, 10% missing values, and a variable FP rate. **F)** 10% missing values, 0.1% FP rate and a variable FN rate. Each combination of error rates and missing values was simulated 20 times.

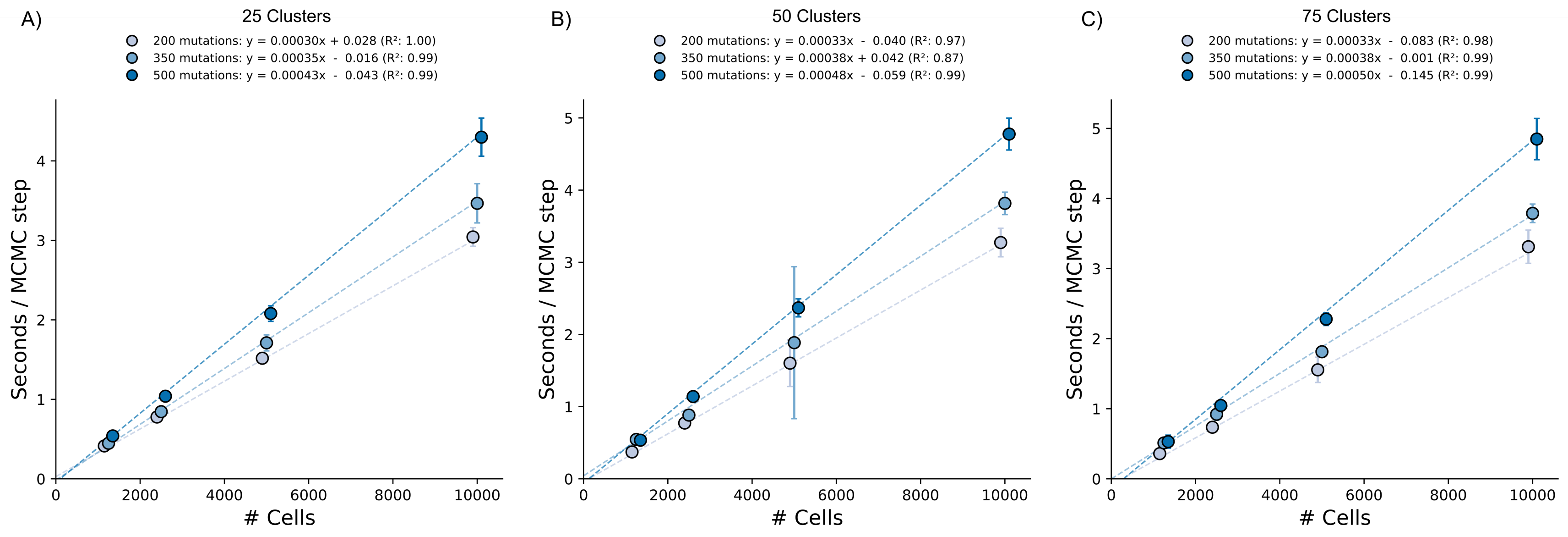


**Figure S9:** Run time per MCMC step for different data sets conforming different number of clusters (25, 50 and 75), mutations (200, 350 and 500) and cells (1,250, 2,500, 5,000 and 10,000). **A)** 25 clusters. **B)** 50 clusters. **C)** 75 clusters Legends display the linear fitted curve and the $R^2$ scores.

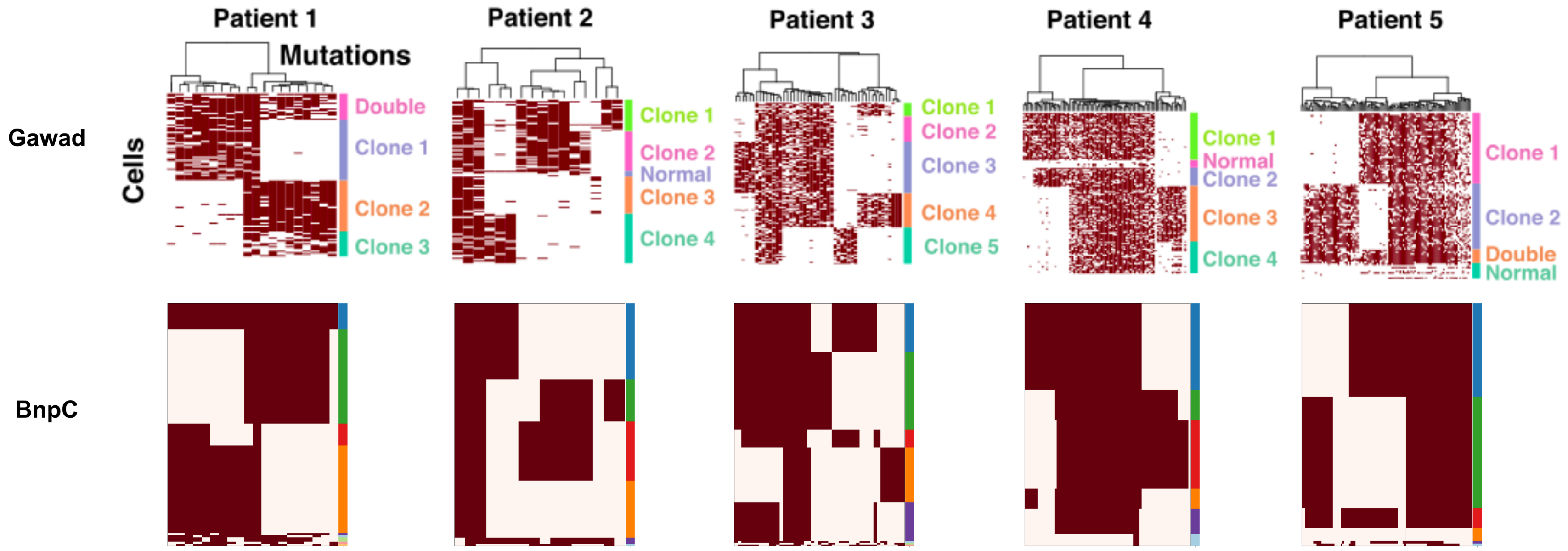


**Figure S10:** Inferred clusters and clonal genotypes by our model for 5 ALL patients compared to Gawad et al. results. Heatmaps depict absence (white) or presence (red) of mutations for every mutation (row) in every cell (column).

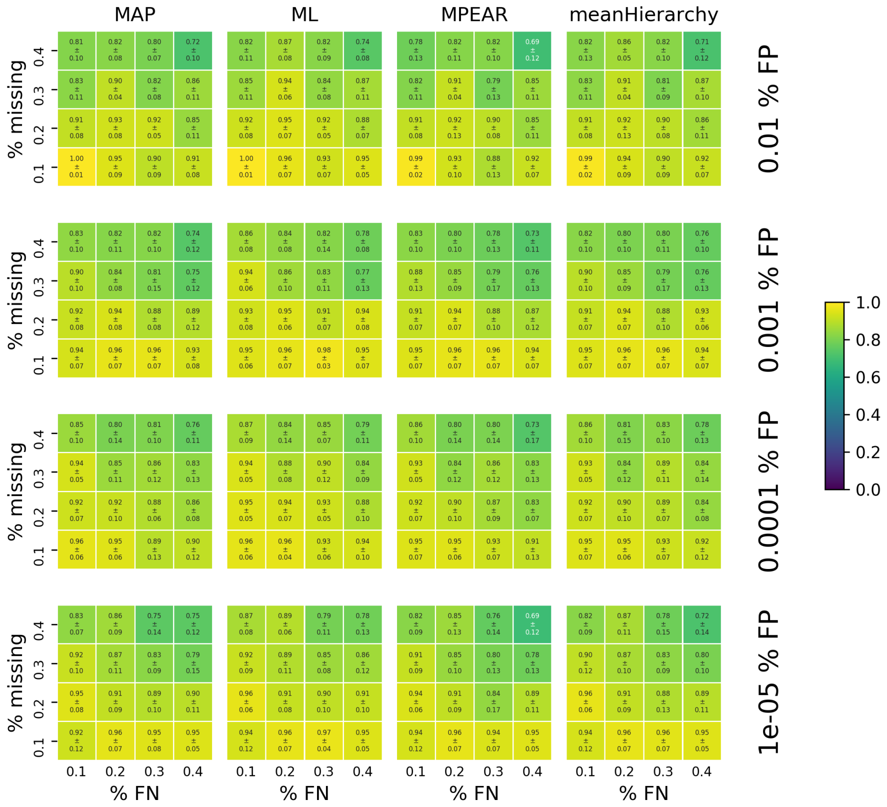


**Figure S11:** Clustering performance measured by the V-measurement of the point estimators MAP and ML, the posterior estimators MPEAR, and our novel estimator named meanHierarchy. The columns represent the different estimators, columns within the heatmaps indicate different false negative rates. The rows represent different false positives rates, rows within the heatmaps indicate a different fraction of missing values. The heatmaps are colored based on the average V-measurement over 10 different trees with an offset of 0.1 and a branching rate of $\frac{1}{3}$. Higher, brighter values indicate better clustering. The ML estimator performs slightly better in most cases but no estimator outperforms the other ones significantly.

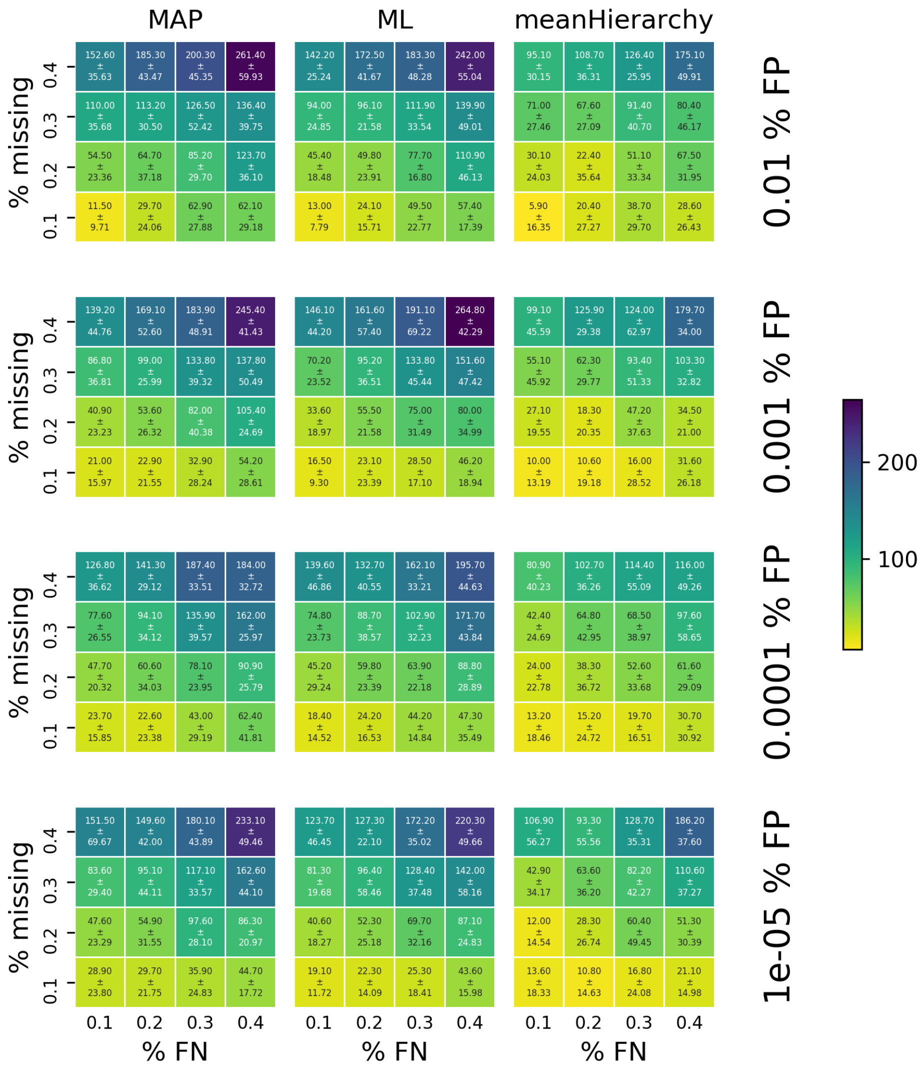


**Figure S12:** Genotyping performance measured by the cell-wise Hamming distance of the point estimators MAP and ML and our novel estimator named meanHierarchy. The columns represent the different estimators, columns within the heatmaps indicate different false negative rates. The rows represent different false positives rates, rows within the heatmaps indicate a different fraction of missing values. The heatmaps are colored based on the average Hamming distance over 10 different trees with an offset of 0.1 and a branching rate of $\frac{1}{3}$. Lower, brighter values indicate better genotyping. Our meanHierarchy estimator outperforms the point estimators in all cases.

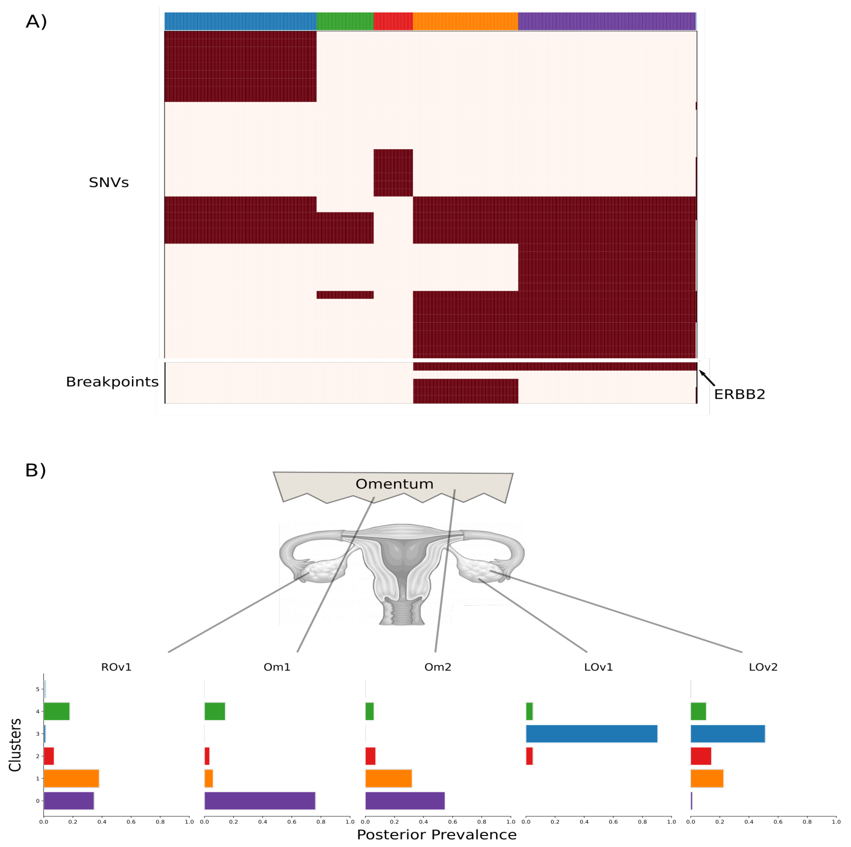


**Figure S13:** **A)** Inferred clones and genotypes by BnpC for patient 9 of the McPherson data set. Heatmap depicts absence (white) or presence (red) of mutations for every mutation (row) in every cell (column). **B)** Estimated prevalence of clones across samples from the posterior distribution estimated by the model.
